# Bees use anthropogenic habitats despite strong natural habitat preferences

**DOI:** 10.1101/278812

**Authors:** Miguel Á. Collado, Daniel Sol, Ignasi Bartomeus

## Abstract

Habitat loss and alteration is widely considered one of the main drivers of the current loss of pollinator diversity. Unfortunately, we still lack a comprehensive analysis of habitat importance, use and preference for major groups of pollinators. Here, we address this gap analysing a large dataset of 15,762 bee specimens (more than 400 species) across northeast USA. We found that natural habitats sustain the highest bee diversity, with many species strongly depending on such habitats. By characterizing habitat use and preference for the 45 most abundant species, we also show that many bee species can use human-altered habitats despite exhibiting strong and clear preferences for forested habitats. However, only a few species appear to do well when the habitat has been drastically modified. We conclude that although altered environments may harbor a substantial number of species, preserving natural areas is still essential to guarantee the conservation of bee biodiversity.

## INTRODUCTION

Pollinators are considered to be of conservation concern worldwide (Goulson et al. 2015). As more than 80% of plants are pollinated by animals (Ollerton et al. 2011), including 75% of crops species (Klein et al. 2007), the extinction of pollinator species is expected to have far-reaching impact on ecosystem functioning and human well-being. For example, in Europe, more than 20% of bees assessed by the IUCN are threatened (Nieto et al. 2014), and in North America there is evidence that the populations of many bee species have drastically declined (Cameron et al. 2011; Bartomeus et al. 2013).

While the current loss of pollinators may have a variety of causes, habitat loss and alteration is widely assumed to be one of the main drivers (Winfree et al. 2011). The conversion of natural habitats into urban and agricultural systems is often drastic and rapid (Frishkoff et al. 2014). Therefore, it is expected that the tolerance limits of many species are exceeded, putting them at risk of extinction. Currently, over 40% of Earth’s terrestrial surface has already been modified by humans (Ellis et al. 2010) and the surface is expected to continue increasing in the next decades (Tilman et al. 2001).

Although human-induced changes in the habitat are expected to have a negative impact on some pollinator species (Winfree et al. 2011), they can also offer ecological opportunities to other species (McFrederick and LeBuhn 2005, Matteson et al. 2008). Pollinators are highly mobile animals, and some are capable of using multiple habitat types (Kremen et al. 2002). For example, some bee species nest in forested habitats while foraging in agricultural habitats (Klein et al. 2003), and some even use highly transformed environments such as those altered by urbanisation and intensive agriculture (Saure 1996, Baldock et al. 2015). If human modified habitats create new opportunities for some pollinators, this may reduce the impact of habitat loss and alteration on pollinator communities and associated ecosystem services (Kremen et al. 2002).

To anticipate the consequences of land use change on pollinator communities, we therefore need to assess the extent to which different species are able to persist in intensively human-modified environments. This will inform us about the sensitivity of species to habitat alteration and will help to identify habitats of conservation importance for protection plans, but unfortunately, we still lack a comprehensive analysis of habitat importance, use and preference for major groups of pollinators. A major obstacle has been the paucity of large-scale datasets on species-habitat associations. In bees, for example, habitat suitability has only been modelled indirectly based on expert knowledge (Lonsdorf et al. 2009, Koh et al. 2015)

In the present study, we use a comprehensive dataset from an extensive monitoring program for bees in the northeast and midwest US to directly estimate habitat use and preference across the entire region. Previous work has established that bee community composition may be strikingly different among habitats (Brosi et al. 2007). For example, in agricultural areas pollinator communities are typically dominated by common species and have few rare or threatened species (Kleijn et al. 2015). Our first goal was to extend this work to ask how bee composition differs among natural and human altered habitats at a regional scale. We assessed habitat importance for bee pollinators, both in terms of species diversity and number of habitat specialists, using data for 15,762 individual bees from 433 species recorded over 15 years. As pollinators are mobile species and the surrounding landscape often determines the presence of a species in the focal habitat (Kremen et al. 2002). we also investigated the effect of the surrounding landscape on determining bee responses.

The assessment of habitat importance can provide important insight into their sensitivity to environmental change. However, the association of a species with a particular habitat does not necessarily indicate that the species is doing well in that habitat, instead, it may simply indicate that it is the only habitat that is available. To better assess species sensitivities, it is necessary to assess habitat preferences, defined as the tendency of a species to be non-randomly associated with certain environments (Rice 1984). Therefore, the second goal of our study was to investigate such species-habitat associations. We used null models to assess habitat preference and avoidance for 45 bee species with a sufficiently large sample of occurrences (species with ≥ 100 independent records). We then characterized their sensitivity to human altered habitats by estimating the extent to which the species occurs in highly-modified environments or, instead, use multiple habitats that buffer them against destruction of their preferred habitat.

## MATERIAL AND METHODS

### Sampling design

Bees were intensively sampled from 2000 to 2015 by USGS Native Bee Inventory and Monitoring Laboratory, their collaborators, and volunteers using pan traps, (∼75%) and hand netting (∼25%). As the sampling was designed to maximize the area covered, not to repeat areas within a season or along years, different locations were selected in each sampling point. Yet, all habitats were sampled enough to cover the entire phenology and with similar monitoring techniques (Figure S1, Table S1). Although sampling was carried out over a larger region, we restricted our analyses to samples taken from the area with the highest sampling effort, covering latitude 35.01 S to 42.79 N and longitude -87.54 W to -69.97 E (Fig. 1).

**Figure 1.**
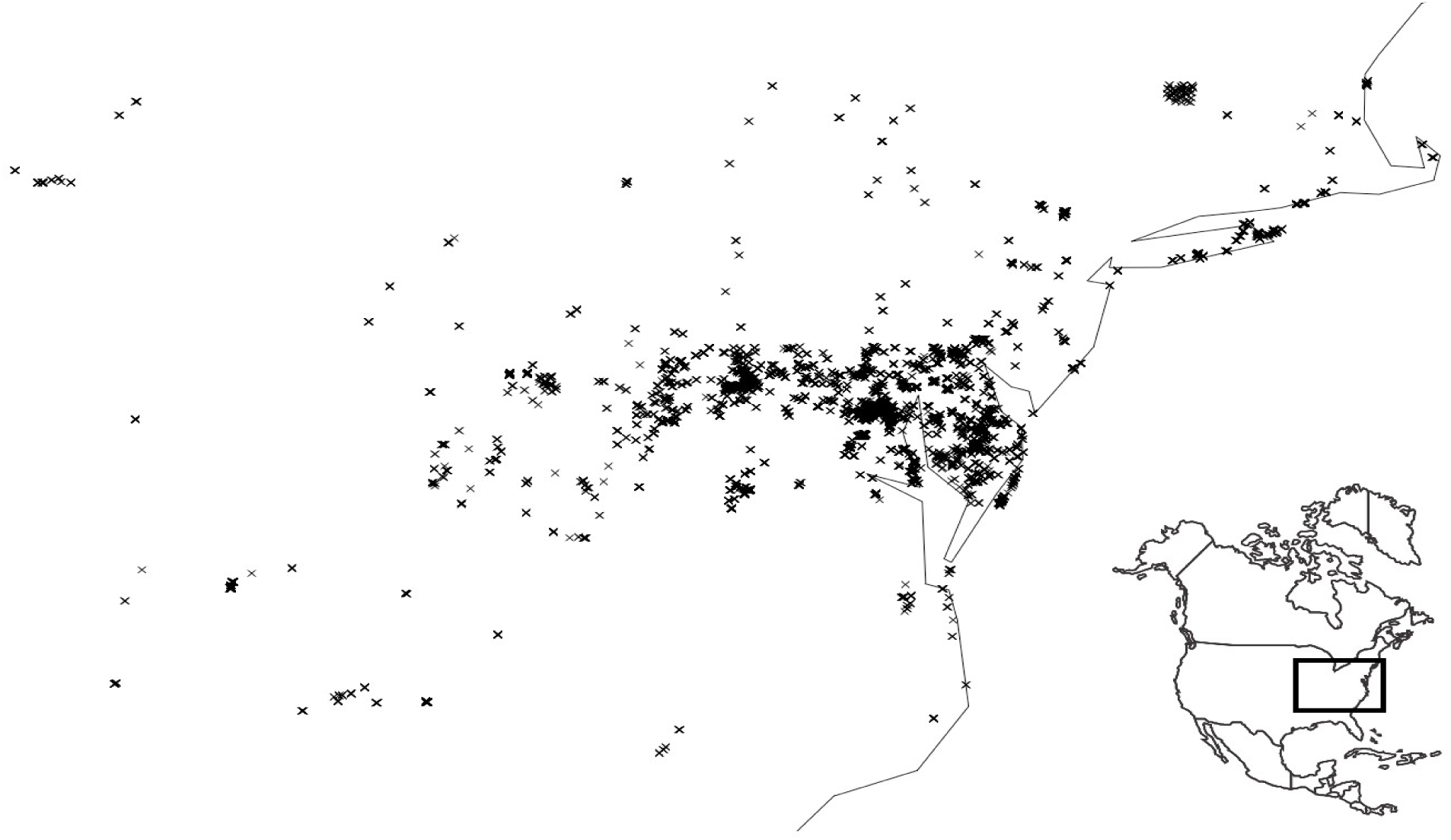
Map of the sampling area. It covers the area from 35.01 S to 42.79 N and -87.54 W to -69.97 E, northeast USA. This area was selected as it represents a large but homogeneous region.

After capture, each specimen was identified to species level by expert taxonomists and the coordinates of the collection site were recorded using GPS. Unidentified individuals or extremely rare species (i.e. those collected only once) were removed from the dataset. Overall, we retained 31,505 individuals, which represent a 66% of the original data. To ensure the independence of the collection events, we excluded from analysis specimens belonging to the same species when collected at the same locality during the same day. After this last filtering, the final dataset comprised 15,762 individuals from 433 species collected from 1,452 different sites, all of which were used in subsequent analysis. All specimens were vouchered at USGS Native Bee Inventory and Monitoring Laboratory.

For each georeferenced sampling site, we extracted habitat information using the National Land Cover Database (NLCD) raster layer (Homer *et al*. 2015) with the R packages *raster, rgdal and stringr* (Bivand, Keitt T, and Rowlingson 2014, Wickham 2015, Hijmans 2015). The 14 habitats considered for this study are described in Table 1. We first extracted the habitat type from the focal point based on the precise coordinates. To take into account the surrounding landscape, we also extracted the habitat composition in a buffer of 1,000 m radius around each focal point; 1,000 m is the maximum distance that most bees under 4 mm of intertegular span can forage (Greenleaf et al. 2007). While our dataset spans 15 different years, information on land cover was only available for 2001, 2006 and 2011. To account for this, bees sampled before 2005 were assigned to habitats based on information from the 2001 layer (28.11 % of our data), those sampled between 2006-2010 were assigned to the 2006 layer (42.32%) and for the rest (2011-2015) we used the 2011 layer (29.57%). To estimate availability of each habitat in our study region, we divided all the pixels of the habitat by the total pixels of the entire study area (Fig. S2 in Supporting information).

**Table 1.**
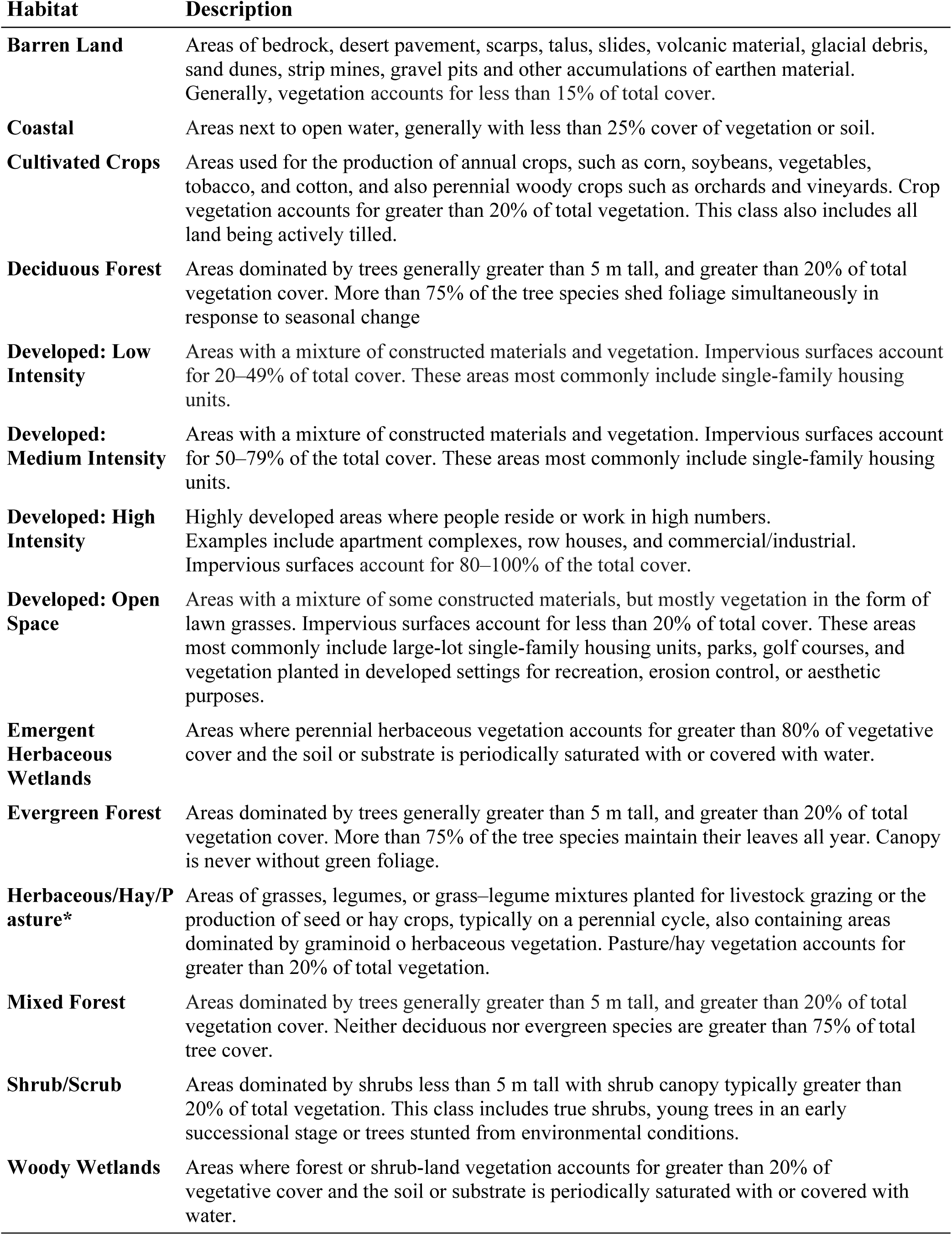
Description of the habitats used to assess importance, use and preference for bee pollinator species, as they are defined and contained in the National Land Cover Database 2011, which is a modified version of the Anderson Land Cover Classification System (J. R. Anderson et al 1976). *Herbaceous and Hay/Pasture are classified as two different habitats in NLCD. We merged them because herbaceous areas in our sampling region are always for livestock (Koh et al 2015).

| Habitat | Description |
| --- | --- |
| **Barren Land** | Areas of bedrock, desert pavement, scarps, talus, slides, volcanic material, glacial debris, sand dunes, strip mines, gravel pits and other accumulations of earthen material. Generally, vegetation accounts for less than 15% of total cover. |
| **Coastal** | Areas next to open water, generally with less than 25% cover of vegetation or soil. |
| **Cultivated Crops** | Areas used for the production of annual crops, such as corn, soybeans, vegetables, tobacco, and cotton, and also perennial woody crops such as orchards and vineyards. Crop vegetation accounts for greater than 20% of total vegetation. This class also includes all land being actively tilled. |
| **Deciduous Forest** | Areas dominated by trees generally greater than 5 m tall, and greater than 20% of total vegetation cover. More than 75% of the tree species shed foliage simultaneously in response to seasonal change |
| **Developed: Low Intensity** | Areas with a mixture of constructed materials and vegetation. Impervious surfaces account for 20–49% of total cover. These areas most commonly include single-family housing units. |
| **Developed: Medium Intensity** | Areas with a mixture of constructed materials and vegetation. Impervious surfaces account for 50–79% of the total cover. These areas most commonly include single-family housing units. |
| **Developed: High Intensity** | Highly developed areas where people reside or work in high numbers. Examples include apartment complexes, row houses, and commercial/industrial. Impervious surfaces account for 80–100% of the total cover. |
| **Developed: Open Space** | Areas with a mixture of some constructed materials, but mostly vegetation in the form of lawn grasses. Impervious surfaces account for less than 20% of total cover. These areas most commonly include large-lot single-family housing units, parks, golf courses, and vegetation planted in developed settings for recreation, erosion control, or aesthetic purposes. |
| **Emergent Herbaceous Wetlands** | Areas where perennial herbaceous vegetation accounts for greater than 80% of vegetative cover and the soil or substrate is periodically saturated with or covered with water. |
| **Evergreen Forest** | Areas dominated by trees generally greater than 5 m tall, and greater than 20% of total vegetation cover. More than 75% of the tree species maintain their leaves all year. Canopy is never without green foliage. |
| **Herbaceous/Hay/Pasture\*** | Areas of grasses, legumes, or grass–legume mixtures planted for livestock grazing or the production of seed or hay crops, typically on a perennial cycle, also containing areas dominated by graminoid or herbaceous vegetation. Pasture/hay vegetation accounts for greater than 20% of total vegetation. |
| **Mixed Forest** | Areas dominated by trees generally greater than 5 m tall, and greater than 20% of total vegetation cover. Neither deciduous nor evergreen species are greater than 75% of total tree cover. |
| **Shrub/Scrub** | Areas dominated by shrubs less than 5 m tall with shrub canopy typically greater than 20% of total vegetation. This class includes true shrubs, young trees in an early successional stage or trees stunted from environmental conditions. |
| **Woody Wetlands** | Areas where forest or shrub-land vegetation accounts for greater than 20% of vegetative cover and the soil or substrate is periodically saturated with or covered with water. |

### Data analysis

#### Habitat importance

We first evaluated the importance of different habitats for bee species using the number of species detected in each habitat (i.e., species richness). Although species richness is a traditional index of habitat importance (Chao and Jost 2012), it treats all species as equal, which may not be optimal for conservation purposes. For instance, a habitat may have high species richness but primarily sustain common species that are widely present elsewhere, whereas another habitat with equal or lower species richness could mostly support rare species that are highly dependent on this particular habitat. Thus, we used in addition to species richness, a metric of habitat strength, as a way of weighting for the species depending on particular habitats (see below).

Habitat strength was calculated using a metric derived from network analysis. The strength of a node (i.e., a single element from a network, in this case the focal habitat) in a bipartite interaction network of species per habitats is defined as the sum of the dependencies of nodes corresponding to the other level in the bipartite network (in this case, the bee pollinator species) linked to that habitat (Bascompte et al. 2006). The dependence of a bee pollinator on a given habitat is calculated as its proportional use of this habitat relative to the other habitats and ranges from zero to one. For example, if a species node has a dependence value near one on a habitat node, we conclude that species depends strictly on that habitat. However, if the dependence is close to zero, the species does not depend on that single habitat and instead, uses other habitats.

To calculate richness and strength for each habitat, we first rarefied each habitat to equalize sampling effort to that of the least sampled habitat. To this purpose, we first calculated the coverage value (the percentage of the total species diversity) for each habitat and then rarefied to the number of individuals necessary for equal coverage of all habitats (Hsieh et al. 2012). The common coverage value used was 0.60, meaning that 60% of species richness from each habitat was sampled to calculate richness and strength. Coverage was calculated using the “iNEXT” package (Hsieh et al. 2012). By using the same coverage for every habitat we avoided that the most sampled habitats were over-represented. However, the total richness at the regional scale (i.e., gamma-diversity) is likely to depend on the area covered by each habitat, independent of the number of samples for each habitat, so we show the proportion of each land cover type (Fig. S2) to aid interpretation of gamma-diversity values.

Finally, we assessed beta-diversity among and within habitats to identify the habitats that are complementary in species composition and determine the degree of species turnover within habitats across space (Whittaker 1960). Among habitat beta-diversity was calculated using Sørensen beta-diversity dissimilarity index across all pairs of habitats (Sørensen 1948, Fig. 2). Habitats were then grouped according to their similarity using a hierarchical cluster analysis. Within habitat beta-diversity was calculated as the slope of the species-samples accumulation curves for each habitat. This metric represents the rate at which new species appear within that habitat as sample size increases (see Fig. 2). The species-samples relationship was almost linear and hence we did not log-transformed the data (Baselga and Orme, 2012 log-transforming the data using natural logarithms produced similar results).

**Figure 2.**
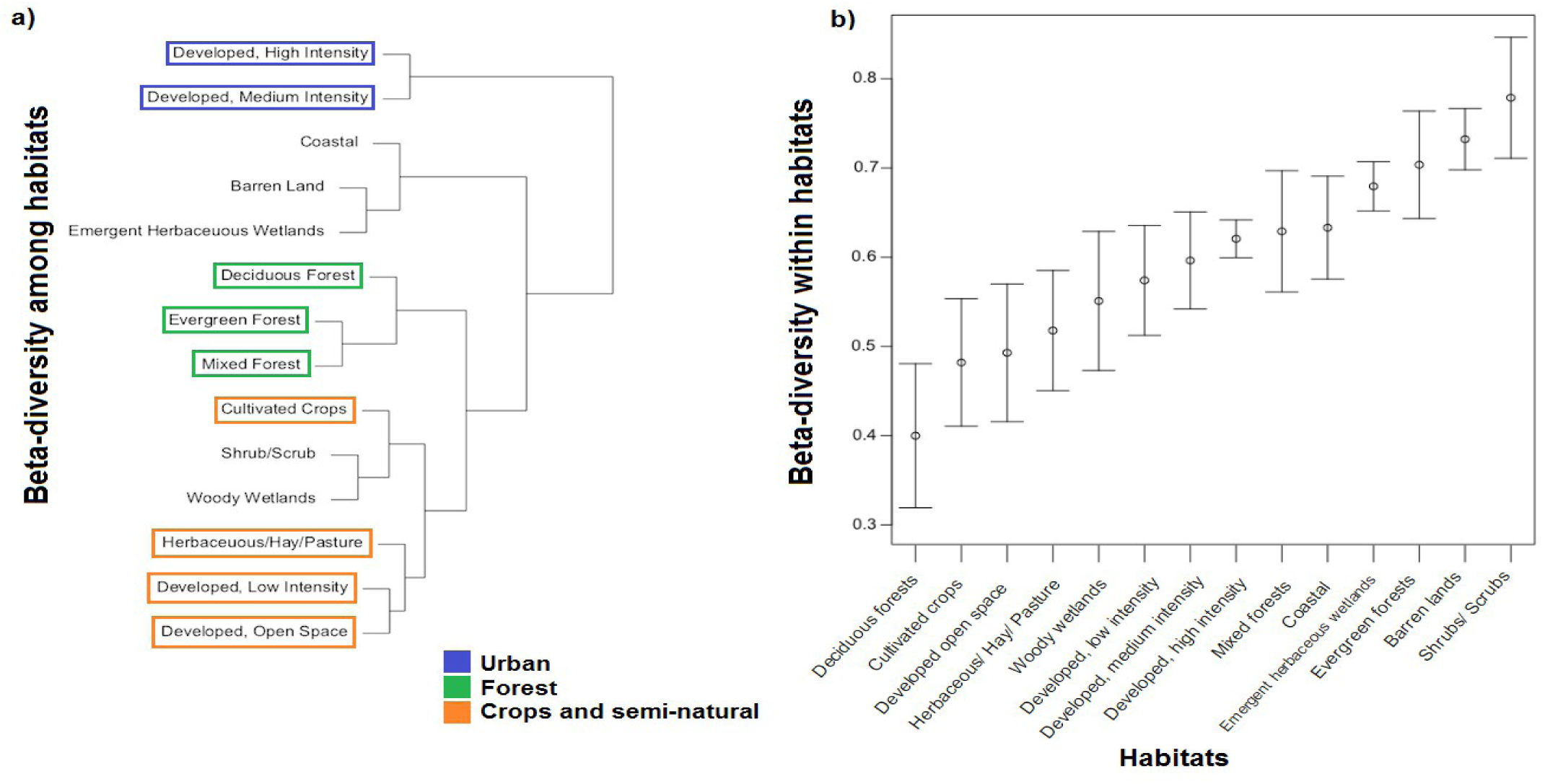
Beta-diversity analysis results: a) using the beta-diversity values among habitat types (Sørensen beta-diversity dissimilarity matrix). We grouped the 14 habitats according to their similarity in community composition. Groups of habitats used for the preference analysis are highlighted. b) Estimated beta-diversity and standard deviation within habitats, calculated as the slopes of the species-sample accumulation curves for each habitat, as an indicator of the rate new species for that habitat appear with increasing sample size. Larger values indicate more rapid gain of new species with increased sample size.

As bees are mobile organisms that likely depend on adjacent habitats in the landscape, we repeated the above analyses at a landscape scale. We classified landscapes at a 1,000 m radius surrounding each sampling site into discrete groups using a k-mean algorithm. The total number of groups (k) was determined using “the elbow method”, where k is the number of clusters beyond which additional clusters no longer improve the model (k = 20, Fig. S3) so 20 types of landscapes were made. These 20 landscape categories range from mainly forested landscapes, to heterogeneous landscapes that include agriculture and forested areas (Table S2).

#### Habitat use and preference

Disentangling habitat use and preference of individual species requires a large sample size for each species studied, thus we only used species that had > 100 independent collection events (n = 45 species). We considered a habitat to be used by a species when at least one individual of that species was sampled in that habitat. To normalize for differences in the number of species occurrences, we did 100 rarefaction events and extracted the means of the number of habitats used by each species.

We defined habitat preferences as the non-random association of a species with certain habitats. Therefore, a species was considered to exhibit habitat preference if it was sampled in a habitat more frequently than expected by chance. Species preferences can be confounded with species distributions if their geographic range only covers some of all available habitats. For example, species distributed only in the northern part of the sampling area may appear to prefer evergreen forests simply because this habitat is more common there. However, this limitation was negligible in our study because the geographic range of the species studied covered the entire study area, implying that all sites could have been potentially occupied by any species if habitat choice was completely random. We compared a habitat-species matrix (i.e., the “observed” matrix) to 1,000 null matrices (i.e., the “expected” matrices). These expected matrices were created by means of the function “nullmodel” contained in the “bipartite” package (Dormann et al. 2009). This function generates random bipartite tables maintaining the sum of rows and columns using Patefield’s algorithm, so the proportional abundance of species and habitats is maintained, but their associations are re-shuffled. We considered that a species exhibited preference for a particular habitat if it was more abundant than the 0.95 quantile of expected abundances. Species less abundant than the 0.05 quantile were considered to avoid that habitat (Sol et al. 2014).

For the sake of clarity, we present in the results section habitat preferences grouped by three main habitat types: 1) urban: developed, high intensity and medium intensity; 2) crops and semi-natural areas: cultivated crops, herbaceous/hay/pasture, developed, low intensity and open space; and 3) forested: deciduous forest, evergreen forest and mixed forest, see Fig. 2 for details. Detailed preferences for each species in each habitat can be found in Table S3.

## RESULTS

### Habitat importance

Our estimates of species richness and strength for each habitat were positively correlated (Fig. 3, Table S3). Despite co-varying positively, the strength values allowed us to differentiate the quality of habitats with similar richness values. For example, except for evergreen forest, which was the habitat with the greatest rarefied species richness (107.7), all rarefied richness values for the three other forested habitats were very similar (range = 91.0 to 91.2 species). Yet, the strength values for these habitats varied substantially, being lowest in mixed forest (22.46), intermediated in woody wetlands (24.56) and highest deciduous forest (25.80). This is because the strength value for a habitat does not only increases with species richness but also when the species are highly dependent on this particular habitat (Fig. S4).

**Figure 3.**
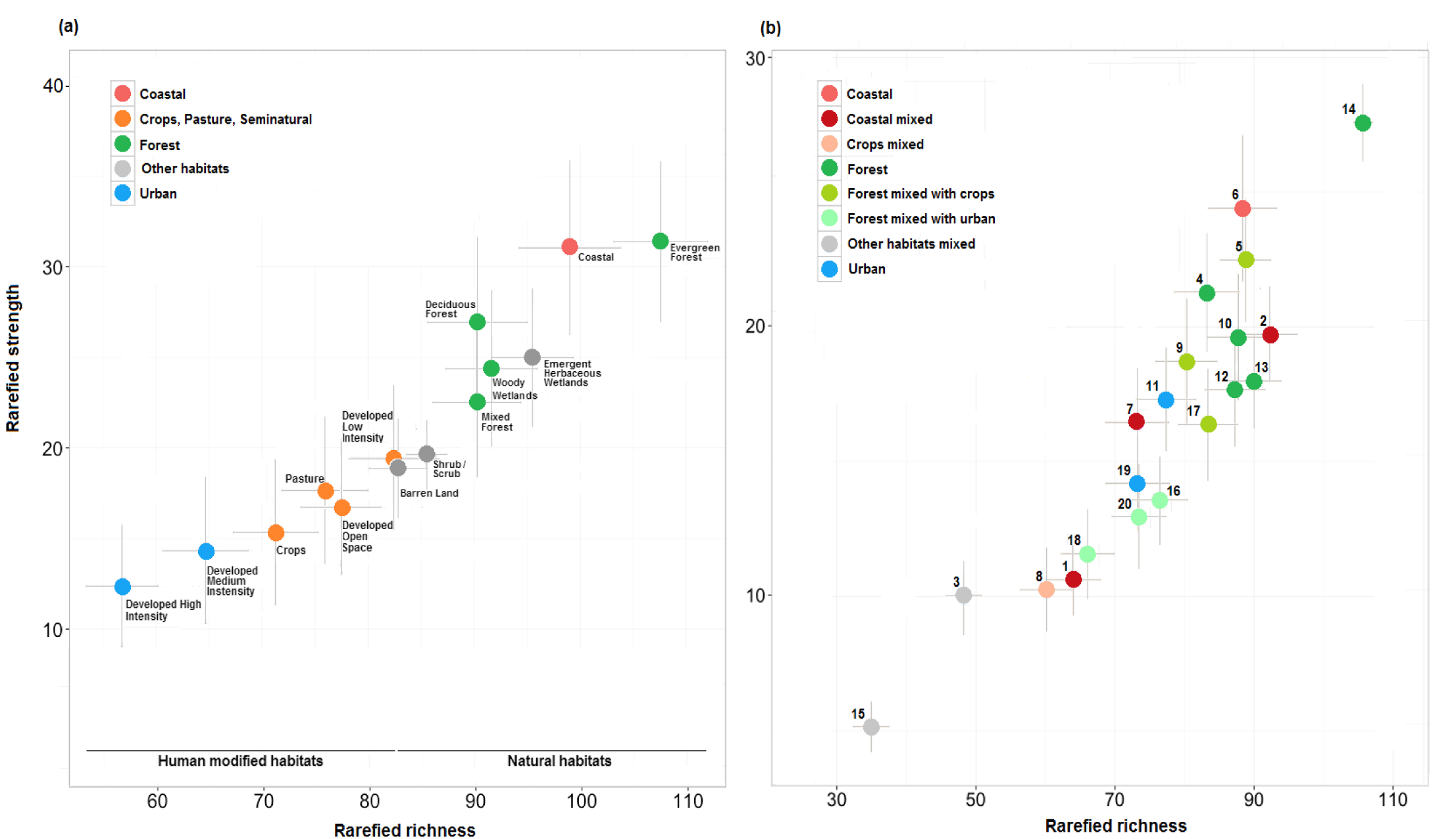
Importance of each analyzed habitat. We defined importance as a function of both strength and richness. Both metrics are correlated, but give different information (see text for details). Each point represents the mean of 100 rarefied strength and richness values for each habitat. Bars are the standard deviation across 100 runs for both strength and richness. a) Shows habitat importance results considering only the habitat where bees were found. While human modified habitats are less important than the natural habitats, they still sustain a substantial amount of pollinator species. b) Shows habitat importance considering landscape composition where species were collected. Similar landscapes were grouped by color; detailed composition of each landscape can be found in Table S1.

**Figure 4.**
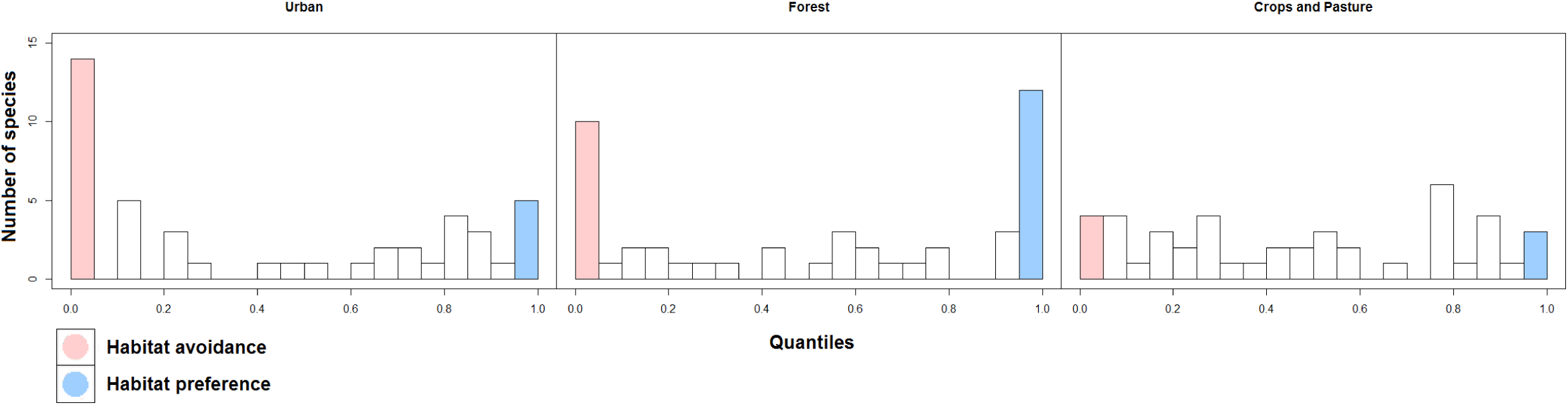
The distribution of species habitat preferences from Table 4. Red bars are the number of species avoiding that habitat, and blue bars are the number of species preferring that habitat. Urban habitats have both avoiders (14 of 45) and exploiters (6 of 45). Forests also have avoiders (10 of 45) but have a higher proportion of exploiters (14 of 45). Crops and pasture are more equally distributed, with few species preferring or avoiding them.

Human-modified habitats sustained a considerable fraction of the regional pool of bee species, however they had lower species richness and strength than less altered and more natural habitats (Fig. 3). However, this lost was in part compensated by the presence of many non-indigenous species. From the 29 exotic bee species we recorded, 22 were collected in urban areas. This is remarkable as only 5% of the sampling area was composed of urban habitat. Exotic species depending on urban habitat were thus contributing to the high habitat strength values reported in urban areas.

The composition of bee species also exhibited remarkable differences across habitats. Cluster analysis on beta-diversity values classified species composition in three main habitat groups: forested habitats (forest), high and medium intensity developed habitats (urban), and crops, pasture and semi-natural habitats (crops and semi-natural) (Fig. 2). The beta-diversity within habitats, which describes the rate at which diversity increases when adding new sampling events, also differed across habitats (Fig. 2). The higher value was observed in evergreen forests, which may explain its high overall species; the lower value instead found in deciduous forests, indicating a high resemblance in species composition across space.

The results at the landscape level highly resembled those at the habitat level. Landscapes dominated by forests had the highest species richness and strength (Fig. 3b, groups 14, 4, 10, 12, 13), even when mixed with crops (groups 5, 9, 17). Coastal areas (6) had also high levels of richness and strength, yet these values diminished again when the proportion of crops and/or pastures increased (1, 3). When crops were the dominant habitat (8), instead, the values of species richness and strength were significantly lower. Finally, urban landscapes (11,19) had low importance than other habitats in terms of species richness, yet their strength values were unexpectedly high because of the presence of urban exploiters (i.e., specialists in developed habitats; Table S2 and Table S3).

### Habitat use and preference

The 45 most common bee species were recorded using most habitats, but 33 out of 45 showed a strong preference for a single habitat (Table S3). After grouping habitats in three main categories (see justification in the methods), 23 out of 45 showed preferences for single groups. Fourteen species preferred natural forested habitats, six preferred urban habitats and three preferred agricultural habitats (Table 2).

**Table 2.** Species habitat preference or avoidance. The first column indicates the number of rarefied habitats used for each species listed, the other three columns show for every habitat (see Fig. 4 for habitat grouping) the habitat preference (> 0.95, marked in blue) or avoidance (> 0.05, marked in red), calculated as the probability of having a higher or lower observed abundance than expected under the null model. * This group of species was merged becasue they are morphologically similar and very difficult to separate by classic taxonomy.

| Species | Habitats used | Pasture and Crops | Forests | Urban |
| --- | --- | --- | --- | --- |
| *Agapostemon virescens* | 12.47 | 0.66 | 0.1 | 0.89 |
| *Andrena carlini* | 11.38 | 0.43 | 0.78 | 0.01 |
| *Andrena cressonii* | 12.41 | 0.53 | 0.61 | 0.02 |
| *Andrena erigeniae* | 11.43 | 0.1 | 1 | 0.01 |
| *Andrena nasonii* | 12.32 | 0.09 | 0.99 | 0.03 |
| *Andrena perplexa* | 10.82 | 0.94 | 0.51 | 0.01 |
| *Andrena violae* | 11.85 | 0.39 | 0.92 | 0.01 |
| *Apis mellifera* | 12.46 | 0.5 | 0.61 | 0.69 |
| *Augochlora pura* | 11.34 | 0.29 | 0.96 | 0.21 |
| *Augochlorella aurata* | 12.3 | 0.05 | 0.75 | 0.5 |
| *Bombus bimaculatus* | 12.04 | 0.05 | 0.99 | 0.04 |
| *Bombus fervidus* | 12.93 | 0.87 | 0.03 | 0.73 |
| *Bombus griseocollis* | 12.28 | 0.19 | 0.43 | 0.82 |
| *Calliopsis andreniformis* | 12.68 | 0.76 | 0.14 | 1 |
| *Ceratina calcarata/dupla/mikmaqi\** | 12.53 | 0.53 | 0.24 | 0.27 |
| *Ceratina strenua* | 12.06 | 0.99 | 0.02 | 0.73 |
| *Halictus confusus* | 12.2 | 0.52 | 0.6 | 0.99 |
| *Halictus ligatus/poeyi* | 12.42 | 0.09 | 0.26 | 0.95 |
| *Halictus rubicundus* | 11.64 | 0.78 | 0.18 | 0.67 |
| *Hylaeus affinis/modestus* | 12.38 | 0.3 | 0.95 | 0.41 |
| *Lasioglossum bruneri* | 11.81 | 0.87 | 0 | 0.96 |
| *Lasioglossum callidum* | 12.35 | 0.99 | 0.01 | 0.11 |
| *Lasioglossum coriaceum* | 12.52 | 0 | 1 | 0.02 |
| *Lasioglossum cressonii* | 11.9 | 0.01 | 1 | 0.04 |
| *Lasioglossum hitchensi* | 12.13 | 0.88 | 0.17 | 0.55 |
| *Lasioglossum illinoense* | 11.59 | 0.79 | 0.77 | 0.89 |
| *Lasioglossum imitatum* | 12.18 | 0.11 | 0.99 | 0.9 |
| *Lasioglossum near_admirandum* | 11.29 | 0.85 | 0.44 | 0.83 |
| *Lasioglossum oblongum* | 12.33 | 0.34 | 0.01 | 0.15 |
| *Lasioglossum pectorale* | 11.59 | 0.58 | 0.11 | 0.15 |
| *Lasioglossum pilosum* | 12.59 | 0.79 | 0 | 0.99 |
| *Lasioglossum tegulare* | 12.51 | 0.76 | 0.01 | 0.25 |
| *Lasioglossum versatum* | 11.96 | 0.5 | 0.7 | 0.25 |
| *Megachile brevis* | 12 | 0.89 | 0.01 | 0.63 |
| *Megachile mendica* | 11.9 | 0.08 | 0.34 | 0.81 |
| *Melissodes bimaculata* | 11.94 | 1 | 0 | 0.84 |
| *Nomada bidentate_group* | 12.52 | 0.42 | 0.98 | 0.02 |
| *Nomada pygmaea* | 11.65 | 0.21 | 0.96 | 0 |
| *Osmia atriventris* | 10.27 | 0.18 | 0.98 | 0 |
| *Osmia bucephala* | 11.94 | 0.29 | 1 | 0.11 |
| *Osmia georgica* | 12.93 | 0.22 | 0.95 | 0.12 |
| *Osmia pumila* | 12.45 | 0.19 | 0.97 | 0.02 |
| *Osmia taurus* | 11.64 | 0.57 | 0.59 | 0.01 |
| *Ptilothrix bombiformis* | 11.53 | 0.78 | 0 | 0.99 |
| *Xylocopa virginica* | 11.65 | 0.28 | 0.6 | 0.79 |

\* This group of species was merged because they are morphologically similar and very difficult to separate by classic taxonomy.

Importantly, species that exhibited a preference for forests also showed a tendency to avoid urban habitats. Thus, 13 of the 14 species that preferred forested habitats avoided urban habitats (Table 2). The only exception was *Lasioglossum imitatum*, who had preference for forested habitats but also presented a high preference for urban habitats (Table 2). Perhaps unexpectedly, some degree of habitat specialization was also detected among urban dwellers (Table 2): seven out of 11 that preferred some type of urbanized environment avoided crops, pastures and forests (Table S3). From all the studied species, the most generalist was the managed bee *Apis mellifera*, which exhibited no preference or avoidance for any habitat.

## DISCUSSION

Four main conclusions can be drawn from our comprehensive analysis of bee species across northeast USA. First, although no habitat appeared to be completely inhospitable to bee pollinators, many species showed a strong preference for natural habitats while consistently avoiding human-modified habitats. Second, the dominant habitat within the landscape was the strongest determinant of species diversity and, contrary to previous studies (Steffan-Dewenter and Tscharntke 1999), heterogeneity only had moderate buffer effects on diversity. Third, and as a consequence of the two previous findings, human-altered environments supported significantly less diversity of species (and had less strength) than the surrounding natural environments. Finally, the loss of biodiversity in human-altered environments could have been higher had not been partially compensated by the addition of human commensals and some exotic species

Natural habitats were the most suitable for bees regarding both importance (richness and strength) and the number of species that preferred them. Evergreen forests in particular exhibited the highest habitat importance in the region, despite harbouring very few habitat specialists. Evergreen forests are widespread in the region and the associated bee species have high spatial beta-diversity, meaning that bee composition largely varies between different evergreen forest sites. These forests comprise flower rich areas like the coastal Pine Barrens that are fragmented and crossed by right of way infrastructures, which can increase their attractiveness for bees (Hill and Bartomeus 2016). Hence, the evergreen forests of our study area may not be representative of more northern evergreen forests. In contrast, bee communities associated with deciduous forests and other natural habitats had lower beta-diversity and lower overall species richness. However, they sustained a large number of habitat specialists, a possibility already advanced in previous studies (Burkle et al. 2013).

As specialised adaptations to particular habitats may limit the success of bee species in other habitats, it is unsurprising that the majority of forest specialist species avoid urban habitats and/or crops and pastures. There is therefore a possibility that many of these species may become extinct, at least locally, if forested habitats disappear from the landscape (Burkle et al. 2013).

Much of the current risk of species loss comes from the replacement of natural forests by crops and pastures, the most frequent alteration of natural habitats (Newbold et al. 2015). Crops and pastures exhibit a significantly low species richness and strength values compared to natural forests (Newbold et al. 2015). Thus, while some species used crops and pastures opportunistically, very few became specialized to exploit them (see also Kleijn et al. 2015). Moreover, within habitat beta-diversity for agricultural habitats was low, reinforcing the view that these habitats sustain a limited set of common habitat generalists. Cropping systems are however highly heterogeneous, ranging from cereal monocultures to diverse flowering cropping systems (Donald 2004) implying that the impact may vary depending on the intensity of the alterations. Although our dataset do not allow for finer scale analyses separating the effects of different crops, current evidence suggests that most modern crop managing practices (i.e., herbicide and insecticide application) are likely to negatively impact on bee populations (Goulson et al. 2015, Woodcock et al. 2016).

While the conversion of natural habitats to cities is not so widespread as the replacement by crops and pastures (Newbold et al. 2015), urbanisation is currently considered one of the most rapid and drastic alterations of natural ecosystems. In line with previous studies (Chapin 1997, Sol et al. 2014), urbanised habitats harboured substantially fewer species than the surrounding natural habitats. The persistence of bee populations in urban habitats may be limited for resource availability. Food resources are often dominated by exotic or ornamental species (Ellis et al. 2012), which few bee species are able to exploit (Bartomeus et al. 2016). However, urban habitats also offer resource opportunities for some species. For example, *P. bombiformis* specializes in exploiting plants from the *Hibiscus* genus, a popular ornamental plant. New opportunities may also emerge for hole nesting bees in the form of human made constructions (Cane et al. 2007). As natural enemies are often scarcer in cities (Sorace and Gustin 2009), these “urban exploiters” may proliferate despite their little opportunity to adapt to the new environments. Likewise, we show that non-indigenous species, proliferate in urbanised environments, being most of the exotic bees collected only in urban areas. Although the presence of urban exploiters and non-indigenous species importantly contributed to increase species richness in urbanised environments, their diversity was low and hence did not fully compensate for the loss of diversity associated with urban avoiders (see also Sol et al. 2017).

Past work suggests that while undisturbed habitats are essential to preserve biodiversity, habitats that have experienced low intensity alterations may still help buffer against extreme diversity loss (Frishkoff et al. 2014, Sol et al. 2017). Our results provide some support to this view, showing that species loss was not as accentuated in moderately modified habitats (Table 2). For example large gardens within cities provide diverse food resources for pollinators, harbouring a higher bee diversity and abundance than city centres. In fact, in Berlin, half of the total German bee fauna was recorded inside the city (Saure, 1996) and in San Francisco, USA, higher mean abundances of *Bombus spp* were found in urban gardens compared with natural parks beyond the city boundaries (McFrederick and LeBuhn 2006). Likewise, some flowering crop fields provide good foraging opportunities for generalist bee species (Magrach et al. 2017), despite low plant diversity and short bloom periods (Donald 2004).

Although the analyses of single focal habitats are essential to establish habitat importance and assess the sensitivity of species to habitat alterations, species diversity typically depends on the mosaic of habitats present in a region (Steffan-Dewenter and Tscharntke 1999). We expected that species able to use multiple habitats would be less vulnerable to habitat modification than species with specific habitat requirements. However, at the landscape level, our results show that the dominant habitat within the landscape was the strongest determinant of species diversity and that heterogeneous landscapes only had intermediate diversity levels. This is exemplified by the finding that forested habitats intermixed with human-modified habitats had lower species diversity than fully forested habitats.

Altogether, our results provide clear evidence that the loss and alteration of natural habitats caused by human activities leads to many “losers” and a few “winners”. Albeit the specific bee-habitat associations vary as a function of the intensity of the alterations and may change in other geographical regions, the pattern we observe might be general (Palma et al. 2017). Admittedly, our estimations of species sensitivity to habitat alterations are conservative, as these analyses were restricted to common species and hence some habitat specialists may have been missed. However, the analyses using the strength index, which did include rare species, consistently showed that the species dependency on a given habitat decreased with the degree of habitat modification (Fig 3). Regardless of any potential bias in the species for which we can assess its preference, our analyses show that the loss of bee diversity could have been higher had not been partially compensated by the colonisation of native opportunists and exotic species (see also Sol et al. 2017). Moreover, the new species may differ from those they replace in functional traits, particularly those that provide environmental tolerance such as life history, body size, multivoltinism and dietary generalism (see Kitahara and Fujii 1994, Bartomeus et al. 2013; Sol et al. 2014; Scheper et al. 2014), and hence probably play different roles in the ecosystem (Bartomeus et al. 2017). As the loss of functional diversity may affect ecosystem functioning and reduce the long-term sustainability of ecosystem services, identifying these functional traits represent an important avenue of future research.

## ACKNOWLEDGEMENT

We thank Rachael Winfree, Montse Vilà, Ferran Sayol, Joan Maspons and Mar Unzeta for discussing the preliminar results from this paper. We thank Sam Droege for the data and bee collection and taxonomical identification and Jamie Stavert for commenting on the final draft.

